# Gene expression profile of the developing endosperm in durum wheat provides insight into starch biosynthesis

**DOI:** 10.1101/2022.10.21.513215

**Authors:** Jiawen Chen, Alexander Watson-Lazowski, Martin Vickers, David Seung

## Abstract

Durum wheat (*Triticum turgidum* subsp. *durum*) is widely grown for pasta production, and more recently, is gaining additional interest due to its resilience to warm, dry climates and its use as an experimental model for wheat research. Like in bread wheat, the starch and protein accumulated in the endosperm during grain development are the primary contributors to the calorific value of durum grains. To enable further research into endosperm development and storage reserve synthesis, we generated a high-quality transcriptomics dataset from developing endosperms of variety Kronos, to complement the extensive mutant resources available for this variety. Endosperms were dissected from grains harvested at eight timepoints during grain development (6 to 30 days post anthesis (dpa)), then RNA sequencing was used to profile the transcriptome at each stage. The largest changes in gene expression profile were observed between the earlier timepoints, prior to 15 dpa. We detected a total of 29,925 genes that were significantly differentially expressed between at least two timepoints, and clustering analysis revealed nine distinct expression patterns. We demonstrate the potential of our dataset to provide new insights into key processes that occur during endosperm development, using starch metabolism as an example. Overall, we provide a valuable resource for studying endosperm development in this increasingly important crop species.

## Background

The endosperm constitutes about 80% of a wheat grain by weight, and is rich in storage reserves that are mobilised during seed germination to nourish the developing embryo [1]. Starch is the main storage carbohydrate in the endosperm, providing much of the calories in wheat-based foods; and storage protein (particularly gluten) is not only important for nutrition, but also for the functional properties of flour for bread and pasta-making [2, 3]. It is therefore important to understand the gene regulation that orchestrates the development of the endosperm and the synthesis of storage reserves.

The endosperm originates from double fertilisation, where a pollen nucleus fuses with the two polar nuclei of the central cell [2, 4]. The triploid nucleus then undergoes numerous mitotic divisions without cell wall formation, forming a multinucleate syncytium, which then becomes cellularised into alveoli by approx. 4 days post anthesis (dpa) [2]. The alveoli cells then differentiate into both aleurone and starchy endosperm cells, and starch synthesis subsequently begins in the latter [5]. The basic structure of the starchy endosperm is formed via further cell divisions and expansion by approx. 10 dpa [2]. It is only after this stage that most of the starch and protein is synthesised: for example, in bread wheat grown in the United Kingdom, grain filling is most active between 14 and 28 dpa [2]. Grain filling is followed by grain maturation and desiccation.

Starch is synthesised in amyloplasts of starchy endosperm cells from sucrose imported into the developing grain from source tissue. It exists as semi-crystalline, insoluble starch granules that are composed of two distinct glucose polymers - amylopectin and amylose. However, wheat (and other species of the Triticeae) is unique among cereals in that it has a “bimodal” distribution of starch granules, where two types of starch granules arise from a distinct temporal pattern of granule initiation [6]. A-type granules are large (18-20 μm in diameter), are flattened in shape, and they start to initiate at approx. 4 dpa and reach their final number by approx. 8 days post anthesis (dpa). B-type granules are small (<10 μm in diameter) and near spherical, and appear at approx. 15-20 dpa [5-7]. The size distribution of A- and B-type granules in wheat has implications on both pasta and breadmaking quality [8-10], and on milling and processing [11-13]. However, we are only beginning to understand the temporal control that coordinates the formation of these two granule types.

While many studies have focused on grain development in hexaploid bread wheat (*Triticum aestivum*), relatively fewer resources exist for tetraploid durum wheat (*Triticum turgidum* ssp. *durum*). Durum wheat is widely grown in semi-arid climates, primarily for pasta production. Worldwide, it represents ≈5% of wheat grown, but it is still the 10^th^ most widely cultivated cereal crop [14]. It is also gaining interest due to its potential for yield and quality improvement [15, 16], its adaptability to warm and dry climates [16-18], and the effects of climate change creating new suitable regions for growing the crop [17]. Further, its tetraploid genome and shorter life cycle compared to bread wheat makes it an ideal experimental model for wheat research. The wheat *in silico* TILLING mutant collection in variety Kronos allows easy isolation of mutants for functional genomics, and mutations in only two homeologs need to be combined for most genes [19, 20]. Durum wheat has now been fully sequenced with a high-quality reference genome [21]. To enable further research into important processes occurring during endosperm development in durum wheat, we generated a high-quality transcriptomics dataset using RNA sequencing of the developing endosperm of variety Kronos, comparable to similar resources available for bread wheat [22]. We demonstrate the potential of our dataset to reveal new insights into grain development, using the process of starch biosynthesis as an example.

## Results and Discussion

### Distinct patterns of gene expression observed during grain development

To explore gene expression patterns that underpin key processes of endosperm development in durum wheat (*Triticum turgidum* ssp. *durum*), we generated a transcriptomics dataset from endosperm tissue extracted throughout grain development, in variety Kronos. Developing grains were harvested from plants grown in controlled environment chambers - starting from the earliest timepoint in which the starchy endosperm could be reliably dissected from the pericarp (6 dpa) and until the grain turned yellow, indicating maturation (30 dpa)(Figure 1A). RNA was extracted from endosperm tissue dissected from the grains (n=3), with each replicate using multiple grains (3-6 depending on the timepoint) harvested from an independent plant. RNA sequencing (RNA-Seq) using Illumina technology yielded between 78,762,223 and 169,888,168 reads per sample. These data were pseudoaligned using *kallisto* [23] to the annotated transcripts from the Svevo v1 durum wheat transcriptome (using Ensembl canonical transcripts only) [21]. A range of 66.356-77.040% of reads per sample were successfully pseudoaligned. We then calculated transcripts per million (TPM) values for each detected transcript (Supplemental File 1). Out of a total of 66,559 annotated canonical transcripts, we detected an average expression over one TPM (for at least one timepoint) for 31,879 transcripts.

**Figure 1:**
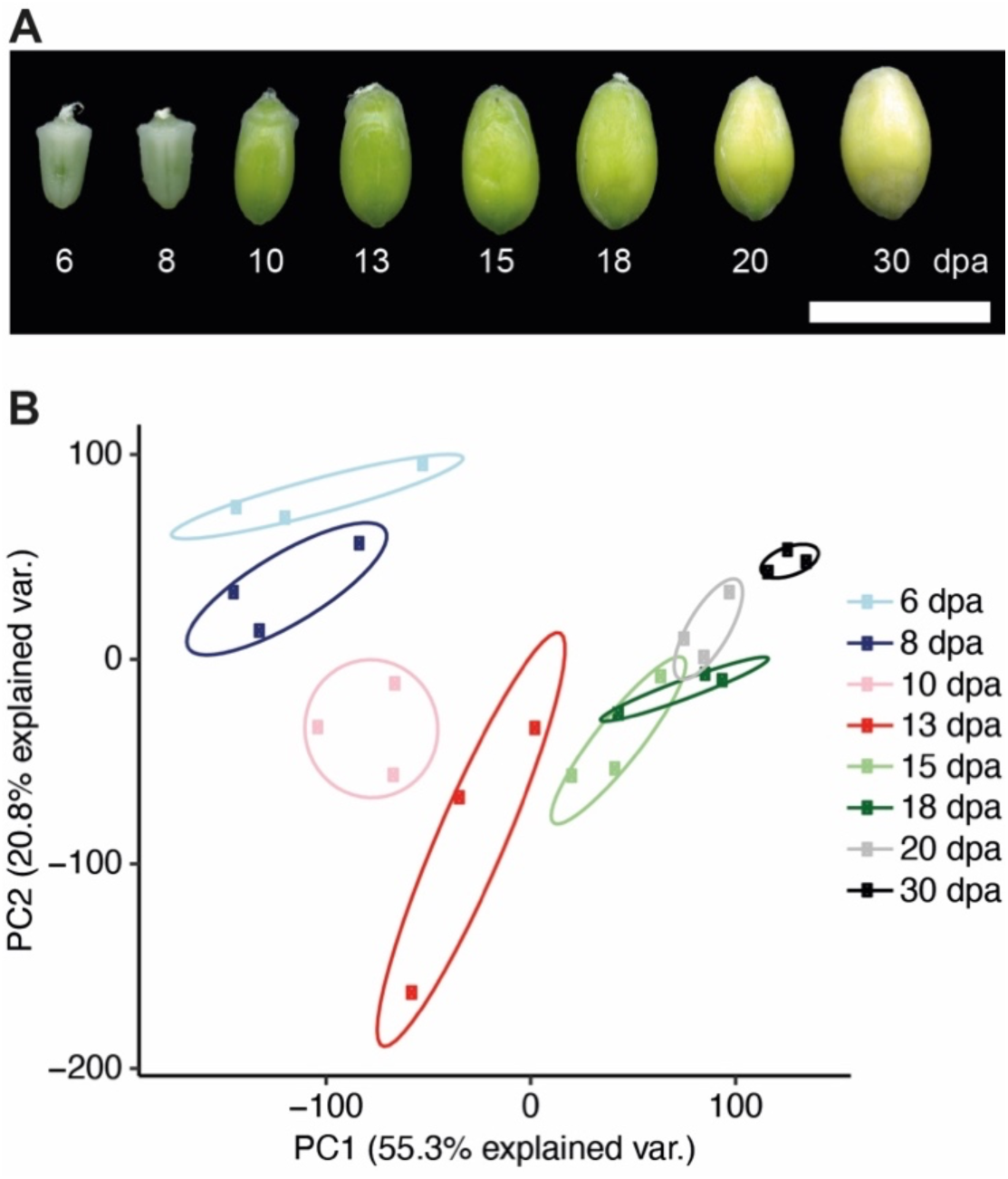
Distinct transcriptome profiles in the endosperm through grain development. **A)** Morphology of grains harvested at each timepoint. Bar = 1 cm, dpa = days post anthesis. **B)** Principal component analysis (PCA) of calculated TPM values. Genes included had a TPM over one for all replicates within our study (15,826 genes included in the analysis). Principal components 1 and 2 (PC1 and PC2) could represent 55.3% and 20.8% of the variation, respectively. Ellipses enclose the three replicates for each timepoint for visualising the spread.

We also calculated TPM values for all annotated transcripts in the Svevo v1 transcriptome, rather than using the Ensembl canonical transcripts only, as a resource for future analysis (Supplemental File 2). These data were not used in the analysis described in the remainder of this paper.

We used a principal component analysis (PCA) to visualise and identify samples that had similar gene expression patterns (Figure 1B). Replicates for each of the early timepoints (6, 8, 10 and 13 dpa) formed distinct clusters that were well separated from each other. However, the later timepoints (15, 18, 20 dpa) appeared to largely cluster together with some overlap between replicates being apparent. This suggests additional factors may contribute to the variation over these timepoints, whereas temporal variation is limited. The latest timepoint (30 dpa) again separated out into a distinct cluster. It is plausible that the largest changes in gene expression are observed during early grain development when the basic endosperm structure is forming [2], and that the expression pattern stabilises as the grain as undergoes filling. The distinct shift in expression pattern at 30 dpa is likely due to processes such as maturation and senescence being initiated. Significantly differentially expressed (DE) genes (DEGs) were calculated using DESeq2 (FDR ≤ 0.05 and log_2_fold change > 1.5) for all pairwise comparisons of the sampled timepoints (Supplemental File 3). This resulted in a total of 29,925 DEGs. When comparing early timepoints (6, 8, 10 and 13 dpa) sequentially, a large number of DEGs were identified (between 2,185 and 6,407 per comparison). However, when comparing within the middle timepoints (15 and 18 dpa) only 34 DEGs were identified. Comparing the last two timepoints (20 and 30 dpa) again showed high numbers of DEGs (5,250). These numbers are consistent with the PCA (Figure 1A) in suggesting that most changes in the endosperm transcriptome happen during the formation of the endosperm rather than during grain filling, with an additional spike as maturation and senescence initiates. All DEGs (genes which were identified as DE in at least one comparison) were then clustered by their TPM values over endosperm development using *Clust* [24], and 14,102 of the DEGs were found to cluster into nine distinct gene expression patterns (Figure 2; Supplemental File 4). Those DEGs that did not cluster did not meet the cut-off criteria of *Clust*, either due to not meeting the minimum TPM, minimum cluster size, or due to having an expression pattern that was perceived by *Clust* as being too flat over time. These nine clusters (cluster C0-C8) will be referred to as the ‘global clusters’ hereafter. Several global clusters represented general up- or down-regulation over endosperm development (C0, C1 and C6). However, the remaining global clusters were more dynamic and may be associated with processes that act during a specific period of development.

**Figure 2:**
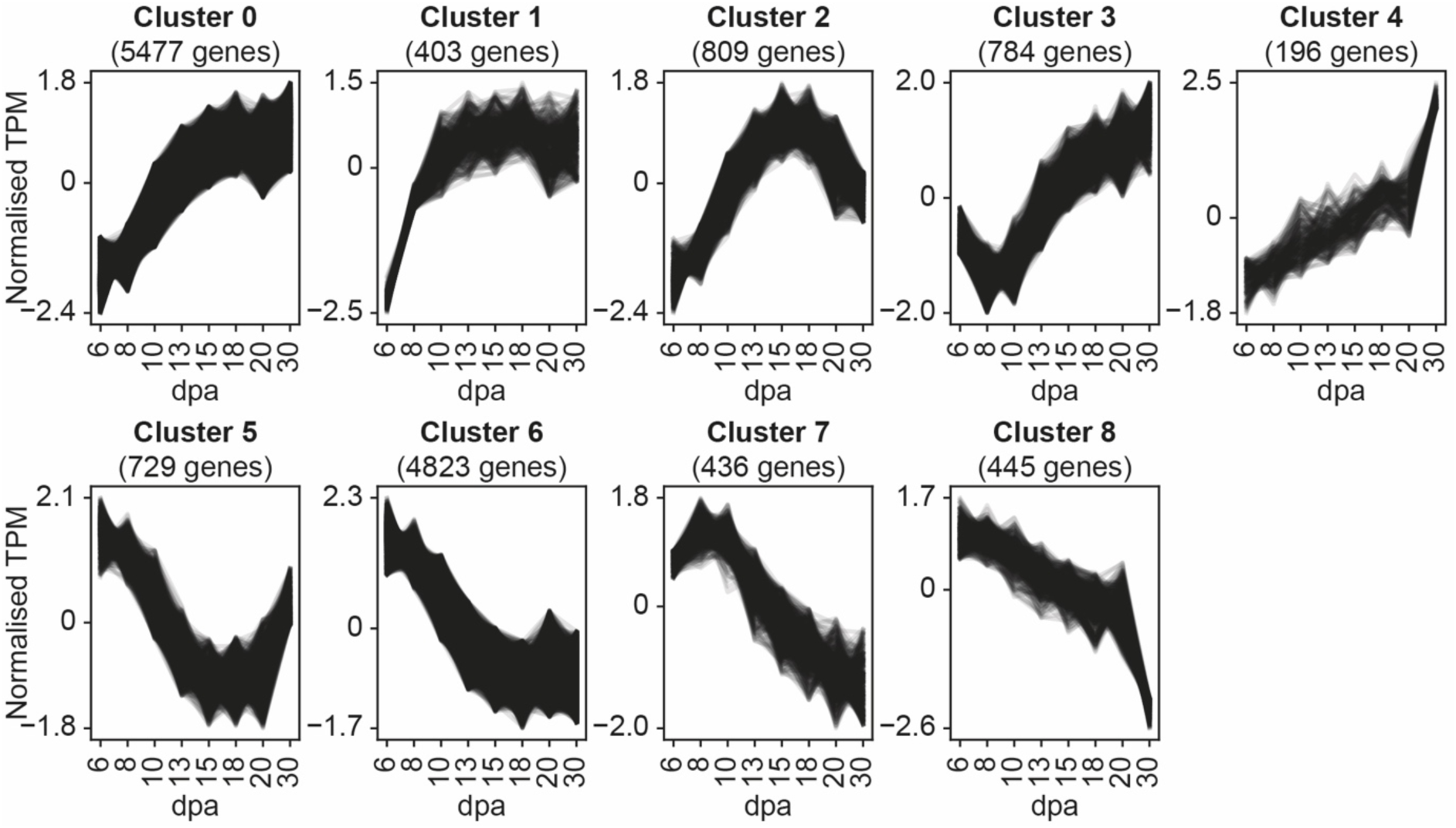
Identification of distinct expression profiles during endosperm development. Out of 29,925 significantly differentially expressed genes (FDR ≤ 0.05 and log_2_ fold change > 1.5), 14,102 clustered into nine distinct expression patterns over endosperm development using *Clust* (tightness value of 5). TPM values were normalised using inbuilt normalisation methods within *Clust*. The number of genes corresponds to the number of genes identified within that cluster. Clustering was performed over endosperm development (days post anthesis (dpa)).

### Analysis of gene expression patterns related to starch metabolism

We analysed the expression patterns of known genes related to starch metabolism, as an example of how a specific process can be studied using our dataset. These genes were subcategorised by their role in starch metabolism: ADP-glucose metabolism, amylopectin synthesis, amylose synthesis, starch phosphorylation, starch degradation, maltooligosaccharide (MOS) metabolism, plastidial sugar transporters and starch granule initiation (Supplemental File 5). Starting from all detected genes (including those that were removed for the DEG/clustering analysis above), we applied a minimum expression filter to remove genes that had very low expression, keeping only those that had an average expression (among the three replicates) of ≥1 TPM for at least one timepoint (Supplemental File 5). Expression patterns for genes within each category were visualised through heatmaps, and hierarchal clustering (HC) was performed to identify genes with similar expression levels and patterns (Figure 3). Almost all starch-related genes that remained after the minimum expression filter had been applied also met the criteria of being a DEG, indicating that they have (to some degree) dynamic expression patterns over development. The implications of this dataset on each category is discussed below.

**Figure 3:**
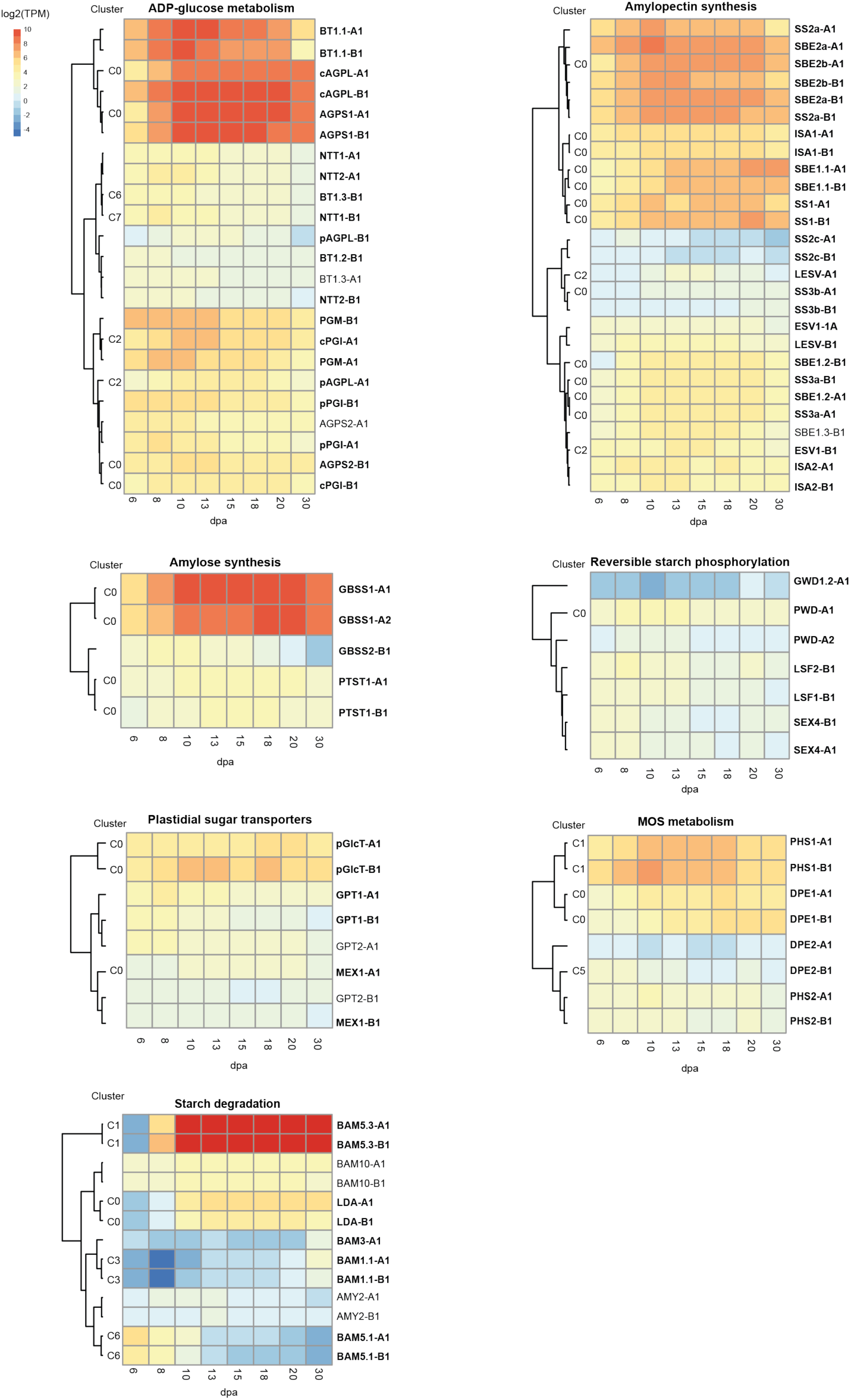
Expression patterns of genes involved in starch metabolism. Average expression over endosperm development and hierarchal clustering of genes within seven categories related to starch metabolism. A total of 91 genes were visualised after filtering out genes with low expression (keeping only genes with average expression of ≥1 TPM for at least one timepoint). Average log_2_ TPM expression values (n=3) were used to illustrate expression, with the same scale used for all categories. Genes in bold were identified as DEGs, and those that were grouped into a global cluster in Figure 2 are annotated with their cluster number (C0-C8). Details of all starch metabolism genes, including average TPM values, can be found in Supplemental File 5. TPM = transcripts per million, dpa = days post anthesis.

### ADP-glucose metabolism

ADP-glucose is the substrate for starch synthesis as it serves as the glucosyl donor for starch synthase-mediated polymer extension. In cereal endosperm, the final step in the conversion of imported sucrose to ADP-glucose takes place in the cytosol and is catalysed by ADP-Glucose Pyrophosphorylase (AGPase)[25, 26]. The ADP-glucose is subsequently imported into the amyloplast by the Brittle1 (BT1) transporter [27]. AGPase is a multimeric enzyme consisting of small (AGPS) and large (AGPL) subunits. AGPS1, which gives rise to the cytosolic small subunit [25, 28], and cAGPL encoding the cytosolic large subunit, are the major subunits expressed in the endosperm – and their expression patterns clustered together with BT1.1. The co-expression of these genes is consistent with these components working together to synthesise and import ADP-glucose into the plastid. The genes encoding plastidial AGPase subunits (AGPS2 and pAGPL) were only lowly expressed (relative to AGPS1 and cAGPL) – suggesting that they do not play a major role in the endosperm. Similarly, other BT1 homologs (BT1.2 and BT1.3), as well as nucleoside triphosphate transporters (NTTs), had low expression compared to BT1.1. The import of ATP into plastids through the NTTs is important for plastidial ADP-glucose synthesis in potato tubers [29] and Arabidopsis guard cells [30], and their low expression in the durum wheat endosperm may reflect the cytosolic synthesis of ADP-glucose in this tissue.

We also made the unexpected observation that wheat has only one set of homeologous phosphoglucomutase (PGM) genes. Both homeologs were expressed in the endosperm, most strongly during early grain development. We ran a phylogenetic tree analysis to better understand the origin of wheat PGM. Closely related barley also had a single PGM gene, which together with the wheat PGMs grouped into the same clade as the cytosolic PGMs of Arabidopsis (PGM2 and PGM3), rather than the clade with the plastidial PGM1 (Supplemental Figure 1). However, both wheat PGM homeologs and most cereal PGM sequences within the PGM2/PGM3 clade were predicted to be plastid localised by TargetP [31]. In the developing bread wheat endosperm, there is measurable PGM activity in both the cytosol and amyloplasts [32], and it remains to be determined how a single PGM paralog can give rise to both activities.

### Amylopectin synthesis

Amylopectin is the primary constituent of starch and is a highly branched polymer with *α*-1,4-linked glucose chains and frequent *α*-1,6-linked branches. Its synthesis requires coordinated action of Starch Synthases (SS) that elongate the polymer chains, Starch Branching Enzymes (SBE) that introduce the branch points, and Debranching enzymes (primarily Isoamylases - ISA) that ensure correct branch pattern formation [33]. Most DEGs within this category that were assigned to a global cluster appeared in C0. Under HC, genes with relatively high expression clustered together – where one branch contained homeologs of SS2a and of the two SBE2 paralogs (SBE2a and SBE2b), which peak at 10 dpa and remain highly expressed until 20 dpa. Their co-expression is interesting because in wheat and maize, these two proteins form a multienzyme complex [34, 35]. The other branch was formed by SS1, SBE1.1 and ISA1 homeologs. SBE1.1 expression levels peaked later than SBE2 (at 20-30 dpa). This is consistent with previous observations in bread wheat endosperm, where SBE2 protein was already visible at 13 dpa and remained present at a similar abundance until 34 dpa, whereas SBE1 was only detectable from 18 dpa and increased in relative abundance towards later grain development [36]. Interestingly, ISA1 and ISA2 did not cluster together under global clustering or HC. In Arabidopsis leaves, ISA1 and ISA2 act exclusively in a heteromeric form [37], but in rice and maize endosperm, the major ISA activity is contributed by ISA1 homomers [38, 39]. The distinct expression patterns between ISA1 and ISA2 may reflect a similar mechanism in durum wheat endosperm. The paralogs of SS2 (SS2b and SS2c) and BE1 (BE1.2 and BE1.3) were detectable but were not as highly expressed as SS2a and BE1.1 in the endosperm. SS3a had higher expression levels than SS3b, but they both had relatively low expression levels compared with SS1 and SS2a.

Homeologs of Early Starvation1 (ESV1) and Like Early Starvation (LESV), ESV1-B and LESV-A, were unique within this category in that they clustered in C2, peaking in expression during mid-grain development (13-15 dpa). ESV1 and LESV were discovered in Arabidopsis and are hypothesised to organise amylopectin within the starch granule [40]. Their expression levels in the durum wheat endosperm were relatively weak compared to the core amylopectin biosynthesis enzymes. However, the recent implication of ESV1 in starch synthesis in the rice endosperm warrants further investigation of these genes in wheat [41].

### Amylose synthesis

Amylose is the minor polymer of starch and consists of *α*-1,4-linked linear chains with very few branches, and is synthesised by a specialised SS isoform, the Granule Bound Starch Synthase (GBSS)[42]. Wheat starch contains approx. 25-29% amylose [43]. All cereals have two isoforms of GBSS – GBSS1 which is mainly expressed in the endosperm and pollen grains, and GBSS2 which is expressed in vegetative organs and the pericarp [44]. In our dataset, we only observed strong expression for GBSS1, which clustered into C0, plateauing in expression towards 18-20 dpa. The very low levels of GBSS2 expression demonstrate that our RNA samples were not substantially contaminated with the surrounding pericarp tissue. Interestingly, we also detected relatively low levels of Protein Targeting to Starch 1 (PTST1) expression. PTST1 in Arabidopsis leaves acts together with GBSS in amylose biosynthesis by facilitating GBSS localisation on starch granules, and *ptst1* mutants produce amylose-free starch [45]. However, in rice endosperm, eliminating PTST1 has only minor reductions on amylose content [46]. The low expression levels of PTST1 in durum wheat makes it unlikely to play a major role in amylose synthesis in the durum wheat endosperm.

### Reversible starch phosphorylation

Cereal starch is almost devoid of covalently bound phosphate [47]. Consistent with this, we observed very low expression levels of the dikinases that phosphorylate starch (GWD and PWD), and the glucan phosphatases that remove phosphates (SEX4 and LSF2). The non-active phosphatase homolog, LSF1, that targets β-amylases to starch granules in Arabidopsis [48], also had very low expression.

### MOS metabolism

Two plastidial enzymes involved in processing MOS were expressed in the endosperm: *α*-glucan phosphorylase (PHS1) and disproportioning enzyme (DPE1) - with PHS1 peaking at around 10 dpa (and appearing in C1) while DPE1 peaked later (18-20 dpa; appearing in C0). PHS1 activity has been observed in the developing wheat endosperm, where it was proposed to participate in starch synthesis either by trimming glucan chains at the granule surface, or by degrading MOS released by ISA during starch synthesis [49]. It is also possible that PHS1 participates in granule initiation. In Arabidopsis, PHS1 is not required for normal starch synthesis or degradation [50], but *phs1* mutations can affect starch granule numbers per chloroplast if other components of MOS metabolism are simultaneously abolished [51]. In rice, the grains of *phs1* knockout mutants are severely shrivelled, particularly when grown at low temperature [52]. The enzyme’s role in granule initiation remains unclear due to the conditional nature of these phenotypes (upon additional mutations and temperature) and should be further investigated in wheat. DPE1 in Arabidopsis leaves participates in starch degradation, where it processes MOS to allow complete degradation to maltose [53]. Since there is very little starch degradation occurring in the developing endosperm, it is not clear what this enzyme does in this tissue. However, there is evidence *in vitro* that the rice PHS1 and DPE1 can physically interact and elongate MOS simultaneously [54].

By contrast to the plastidial PHS1 and DPE1, their cytosolic counterparts, DPE2 and PHS2, showed very low expression. It is plausible that the activity of these enzymes is not important in the developing endosperm, given that cytosolic MOS processing is typically associated with the further metabolism of maltose generated during starch degradation in leaves [55, 56].

### Plastidial sugar transporters

We observed very low expression of the plastidial maltose exporter, MEX1, consistent with there being low starch degradation to generate maltose for cytosolic processing. Interestingly, we observed substantial expression of the glucose transporter pGlcT. In Arabidopsis leaves, pGlcT works together with MEX1 to export products of starch degradation [57]. Its role beyond Arabidopsis leaves remains to be determined. Also, we observed generally low expression for all glucose-6-phosphate (Glc6P)/phosphate translocators (GPT), which are important in heterotrophic or part-heterotrophic tissues such as the embryo and guard cells of Arabidopsis [58, 59] and rice pollen [60], where they import glucose-6-phosphate (Glc-6P) into the plastid to support metabolism, including ADP-glucose synthesis. Their low expression levels are aligned with the cytosolic synthesis of ADP-Glucose in wheat endosperm.

### Starch degradation

The low expression of most starch-degrading enzymes is consistent with the endosperm being primarily a site of starch synthesis. The major plastidial α-amylase and isoamylase activities, AMY3 and ISA3, were filtered out due to very low expression at all timepoints (Supplemental File 5). Expression of most β-amylase isoforms was generally low, except the two homeologs of BAM5.3 (encoded on chromosomes 5A and 4B), which had extremely high levels of expression during grain development from 10 dpa onwards (and appeared in C1). Interestingly, BAM5.3 is an ortholog of Arabidopsis BAM5 and BAM6, and is localised to the cytosol [61]. The role of β-amylases is not understood in cereal grains as even during grain germination, starch degradation is primarily carried out by the AMY1 family of α-amylases [62, 63]. It has even been postulated that the β-amylase accumulated during grain filling may act as a storage protein [64]. BAM5.3 appears to be the primary isoform accumulated during grain development in durum wheat, and its function in the cytosol deserves further investigation. Notably, we also observed substantial expression of limit dextrinase (LDA), which in barley has been proposed to be involved starch granule initiation, since a mutant in LDA has altered distribution of A- and B-type granules [65]. It is possible that it may have a similar role in starch biosynthesis in wheat.

Although the AMY1 family of α-amylases are widely known to be involved in starch degradation during germination, we examined the expression patterns of AMY1 homologs in the developing durum wheat endosperm. This family is characterised by having an N-terminal signal peptide, which allows enzymes synthesised *de novo* to be secreted into the endosperm from the aleurone during germination [63]. We identified 15 AMY1 genes in durum wheat. Surprisingly, one homolog of AMY1 (encoded by homeologs TRITD5Av1G228350 and TRITD5Bv1G227200) was strongly expressed in the endosperm during early grain development, peaking at 8 dpa, before decreasing to almost undetectable levels after 15 dpa (Figure 4A). The role of this secreted *α*-amylase in the endosperm during grain development is yet to be investigated. The other AMY1 homologs, as expected, had very low or almost no expression in the developing endosperm (Figure 4B).

**Figure 4:**
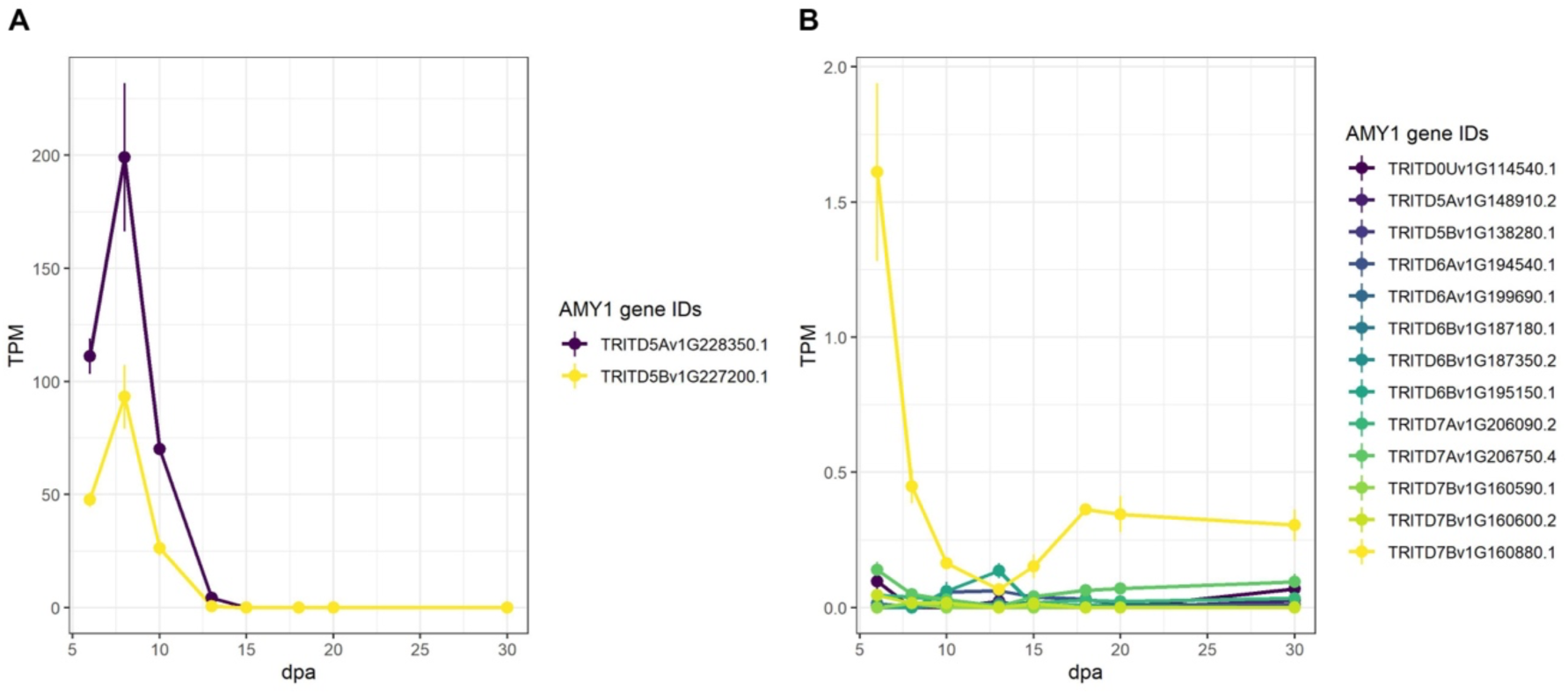
Expression patterns of AMY1 genes in developing wheat endosperm. Transcript accession numbers are the canonical transcripts from Ensembl Plants. **A)** Average TPM values of the two homeologs with high expression in the endosperm. Values are the mean ± SEM of the *n* = 3 replicates. **B)** As for A, but for AMY1 genes with low expression in the endosperm.

### Granule initiation

A distinct set of proteins involved in starch granule initiation in Arabidopsis has recently been discovered. However, research investigating their roles in mediating the distinct spatiotemporal pattern of A- and B-type granule initiation in wheat is still on-going [66, 67]. The genes encoding these proteins showed distinct expression patterns that are consistent with their proposed roles in A- and B-type granule initiation (Figure 5A). Starch Synthase 4 (SS4) was primarily expressed during early grain development, consistent with a role in establishing correct granule number per amyloplast in early grain development [68]. B-Granule Content1 (BGC1) not only limits granule initiations to a single A-type granule per amyloplast during early grain development, but it also has a critical role in promoting B-type granule initiation during later grain development [6]. Consistent with this, BGC1 expression increases during grain development, and is highly expressed at 15-20 dpa when B-type granules initiate. For both SS4 and BGC1, the patterns of transcript accumulation are consistent with patterns of protein accumulation observed through immunoblotting [68]. Myosin Resembling Chloroplast Protein (MRC) was expressed primarily during early grain development, showing essentially an inverse expression pattern to BGC1. This distinct expression pattern appeared in C6, one of the few starch-related genes to do so (alongside BAM5.1). In an accompanying paper, we demonstrate that MRC is a repressor of B-type granule initiation specifically during early grain development, as mutants of durum wheat defective in MRC show the early initiation of B-type granules [69].

**Figure 5:**
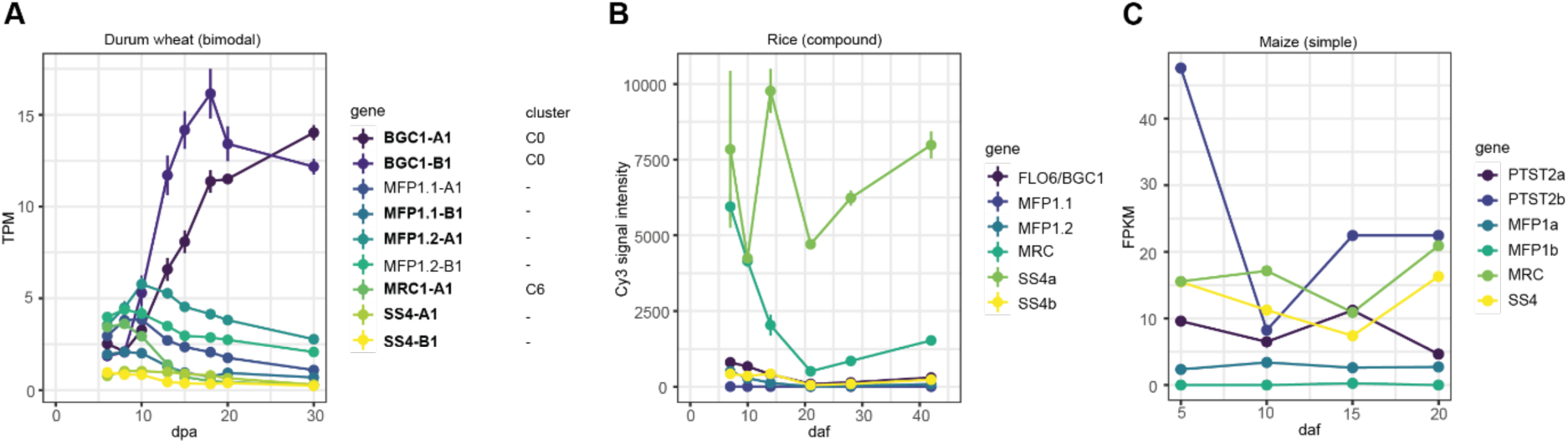
Expression patterns of starch granule initiation genes in three species with different endosperm starch granule morphology. **A)** Average TPM values (n = 3) of starch granule initiation genes in durum wheat endosperm. Genes in bold were identified as DEGs, and numbered cluster annotations (beneath ‘cluster’) refer to the corresponding global cluster from Figure 2 if the gene was included. **B)** Average expression values (n=3) from microarray analysis of starch granule initiation genes from rice developing endosperm, extracted from a public database (https://ricexpro.dna.affrc.go.jp/) [71]. FLO6/BGC1 = LOC_Os03g48170, MFP1.1 = LOC_Os05g08790, MFP1.2 = LOC_Os01g08510, MRC = LOC_Os02g09340, SS4a = LOC_Os01g52260, SS4b = LOC_Os05g45720 **C)** Expression values (n=3) from RNA sequencing analysis of starch granule initiation genes from maize developing endosperm, using data from Qu *et al*. 2016 [72]. PTST2a = GRMZM2G104501, PTST2b = GRMZM2G122274, MFP1a = GRMZM2G106233, MFP1b = GRMZM2G142413, MRC = GRMZM2G104357, SS4 = GRMZM2G044744. For A and B, error bars represent the standard error of the mean. For C, no data was available on the error values. dpa = days post anthesis. daf = days after flowering.

So far, the Mar1 Filament-like Protein (MFP1) is the only known granule initiation protein identified in Arabidopsis that has not been explored in wheat. Interestingly, all cereals have two paralogs of MFP1 (MFP1.1 and MFP1.2)[70], which were both expressed in the durum endosperm (Figure 5A). Given that MFP1 is a thylakoid-associated protein in Arabidopsis leaves [70], the expression of this protein in the endosperm (where amyloplasts lack true thylakoid membranes) is noteworthy, and its localisation and role in the endosperm should be investigated.

To assess how specific these expression patterns are to species that have A- and B-type granules, we examined the expression patterns of these genes in the endosperm of rice and maize using publicly available data. Rice produces compound granules, where granules are initiated during early grain development and there is no second wave of granule initiation in later grain development that leads to B-type granules; and maize produces simple granules without distinct populations of different granule sizes [67]. Among the granule initiation genes, BGC1 had the most different expression patterns when comparing durum wheat with rice and maize. Rather than peaking during mid-late grain development, BGC1/Floury Endosperm 6 (FLO6) expression in rice was highest at the beginning of endosperm development and then decreased, increasing again only slightly in late grain development (after 20 dpa)(Figure 5B). PTST2b (PTST2 is the ortholog of BGC1) in maize was also high during early grain development, dipping and then plateauing in later grain development (Figure 5C). It is therefore tempting to speculate that change in the BGC1 expression pattern is one of the key factors that facilitates the formation of A- and B-type granules.

### Future outlook

We have generated a high-quality transcriptomics dataset for endosperm development in durum wheat, and we demonstrated its potential to provide new insights and hypotheses into key processes during grain development, using starch metabolism as an example. Other processes to be investigated may include protein and lipid biosynthesis, cell wall biosynthesis, micronutrient transport, as well as development of endosperm structure and subsequent maturation in general.

## Materials and Methods

### Plant growth and harvest

Wheat plants were grown in controlled environment rooms (CERs) at 60% relative humidity with 16 h light at 20°C and 8 h dark at 16°C. The CER light intensity was 400 μmol photons m^−2^ s^−1^. Plants were marked for anthesis when an anther was visible from the spike, and this was taken as day 0. Checking for anthesis and harvesting were performed at 1 – 2 hours after midday. Four spikes from each individual plant were used, each for a different developmental time point. Three biological replicates per time point were collected from independent plants. Therefore, some of the biological replicates for separate time points were spread over several of the same plants. For sample collection, the whole spike was cut at the base, frozen in liquid nitrogen and stored at -80°C. Then, grains were threshed from the spikes on dry ice, avoiding the middle grain of each spikelet, and stored at -80°C again. Endosperms were subsequently dissected on dry ice, frozen in liquid nitrogen and stored at -80°C until RNA extraction.

### RNA extraction and sequencing

Total RNA was extracted (n=3) from the frozen endosperm tissue using the RNeasy PowerPlant kit (Qiagen). An on-column DNase digest was incorporated during the extraction. Either 6 (for 6, 8 and 10 dpa), 4 (for 13, 15, and 18 dpa) or 3 (for 20 and 30 dpa) endosperms were pooled from each spike for each RNA extraction. Quality control, poly-A selection library preparation and RNA sequencing (RNA-Seq) using a NovaSeq 6000 machine (Illumina) was carried out at Novogene (Cambridge, UK). Each sample returned between 78,762,223 and 169,888,168 million clean 150 base pair paired end reads. These clean RNA-Seq reads can be downloaded from the NCBI Gene Expression Omnibus repository (Accession: GSE216253).

### TPM value calculation and differential expression

Clean reads from above were pseudoaligned to the Svevo v1 Durum wheat transcriptome downloaded from Ensembl Plants, either using canonical transcripts only (Supplemental File 1) or all transcripts (Supplemental File 2), using *kallisto* [23]. The default settings were used for mapping. A range of 66.356-77.040% of reads per sample aligned to the transcriptome. Both estimated count values and normalised expression as transcript per million (TPM) for each gene were retrieved from the pseudoalignment. *DESeq2* [73], run though the *DEapp* shiny [74], was then used to identify significantly differentially expressed genes (DEGs; significance cut-offs used - FDR ≤ 0.05, log_2_ fold change > 1.5 and a minimum count cut-off of 10 counts in at least three replicates).

### Global clustering analysis

Clustering analyses were performed using *Clust* [24], on all of the DEGs identified. The DEGs were first filtered by setting a minimum threshold of 10 TPM for the sum of the TPM values of the three biological replicates, in at least one time point. The DEG TPM values (as separate biological replicates) were then used as the input for the analysis, which implemented the suggested in-built normalisation methods of *Clust* (log_2_, Z-score and quartile normalisation) [24]. Parameters used included a tightness value of 5, a minimum cluster size >10, and the removal of clusters with flat expression profiles over time. Several tightness values were tested, and the number of clusters identified plateaued at a tightness value of 5.

### Principal component analysis and data visualisation

Principal component analysis (PCA) was carried out in R first using the ‘prcomp’ function from the ‘stats’ package to generate the components of the PCA, then plotted using the ‘ggbiplot’ function from the ‘ggbiplot’ package. TPM values were used as the input for the PCA, with genes with a TPM over one for all replicates being included. For heatmap visualisation and hierarchal clustering, the R function ‘pheatmap’ from the ‘pheatmap’ package was used. Only the rows (genes) were clustered and not the columns (timepoints), otherwise default settings were used. Average log_2_ TPM values for the three replicates at each time point were used as the input for heatmap and hierarchal clustering analyses. Analyses were carried out in version 1.0.12 of the pheatmap package, running on R version 4 (https://cran.rstudio.com/web/packages/pheatmap/index.html).

### Identification of durum wheat starch genes

Genes related to starch synthesis in wheat were manually curated: Orthologs in durum wheat were identified using BLASTp against the Svevo v1 Durum wheat proteome on Ensembl Plants [75], as well as using the “Orthologues” tool Ensembl Plants. In both cases, a known homolog from Arabidopsis, bread wheat, rice, or other species was used as a query sequence. The full list of genes analysed can be found in Supplemental File 5.

## Declarations

### Availability of data and materials

The full dataset generated and analysed during the current study is available in the NCBI Gene Expression Omnibus repository (https://www.ncbi.nlm.nih.gov/geo/; Accession: GSE216253).

### Authors’ contributions

J.C and A.W-L conceived and designed the study, conducted RNA sample preparation, analysed RNAseq data and significantly contributed to writing the manuscript. M.V designed bioinformatics analyses and analysed RNAseq data. D.S conceived and designed the study and significantly contributed to writing the manuscript. All authors read and approved the final manuscript.

## Supplemental Figures and Files

**Supplemental Figure 1**. Phylogenetic tree of PGM isoforms.

**Supplemental File 1**. Calculated TPM values from RNAseq reads aligned to the canonical transcripts of the Svevo v1 durum wheat transcriptome.

**Supplemental File 2**. Calculated TPM values from RNAseq reads aligned to all transcripts of the Svevo v1 durum wheat transcriptome.

**Supplemental File 3**. DEGs of pairwise comparisons between endosperm developmental time points.

**Supplemental File 4**. DEGs in each global cluster identified during endosperm development.

**Supplemental File 5**. Expression pattern of wheat starch metabolism genes.

