## Supplemental Figure 1 for "Gene expression profile of the developing endosperm in durum wheat provides insight into starch biosynthesis"

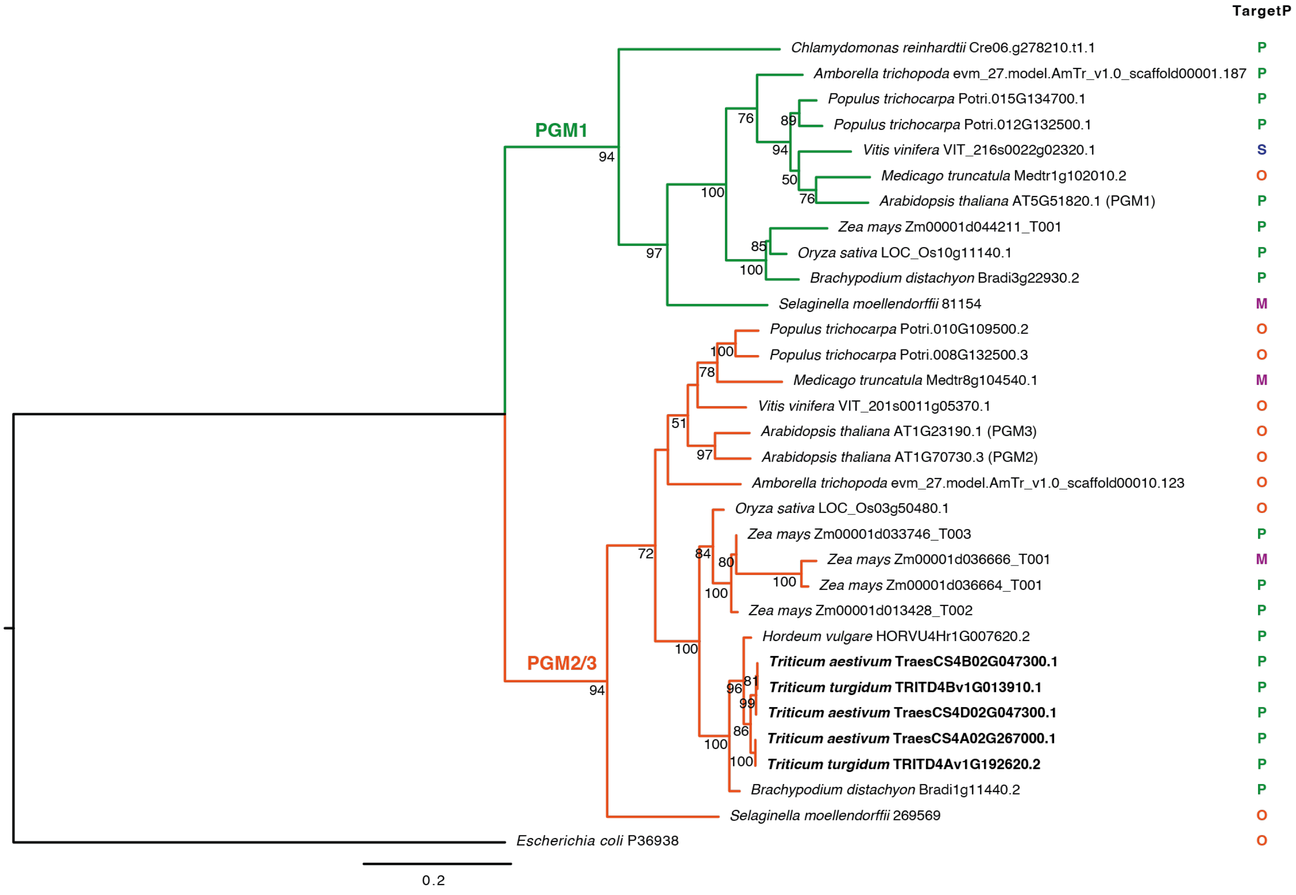


**Supplemental Figure 1: Phylogenetic tree of PGM isoforms.** Sequences of PGM isoforms were obtained from Phytozome v13 (Goodstein et al., 2012; doi: 10.1093/nar/gkr944). Sequences were aligned using MAFFT (Rozewicki et al., 2019; doi: 10.1093/nar/gkz342) and a maximum likelihood tree was constructed using RAxML (Stamatakis 2014; doi: 10.1093/bioinformatics/btu033) with 1,000 bootstrap replicates. Bootstrap values greater than 50 are shown next to the node. Branch lengths represent the number of substitutions per site, indicated by the scale bar. TargetP (Emanuelsson et al., 2007; doi: 10.1038/nprot.2007.131) was used to predict the subcellular targeting of each isoform, either as plastidial (P), mitochondrial (M), secreted (S) or other (O).
